# Migrasome formation is initiated preferentially in tubular junctions by alternations of membrane tension or intracellular pressure

**DOI:** 10.1101/2023.08.03.551756

**Authors:** Ben Zucker, Raviv Dharan, Dongju Wang, Li Yu, Raya Sorkin, Michael M. Kozlov

## Abstract

Migrasomes, the transient vesicle-like cellular organelles, arise on the retraction fibers (RFs), the branched tubular extensions of the plasma membrane generated during cell migration. Migrasomes form in two steps: a local RF swelling is followed by a protein-dependent stabilization of the emerging spherical bulge. Here we approached experimentally and theoretically the previously unaddressed mechanism of the initial RF swelling. We hypothesized that the swelling can be driven by alterations of the generic mechanical factors, the RF’s luminal pressure and membrane tension. To examine the effects of pressure, we exposed migrating RF-producing cells to a hypotonic medium and observed the formation of migrasome-like bulges with a preferential location in the RF branching sites. To test the results of tension variations, we developed a biomimetic system of three membrane tubules connected by a junction and subjected to controlled membrane tension. An abrupt increase of tension resulted in a migrasome-like bulge formation in the junction and in the tubular regions. Following the formation, the tubule’s bulges moved toward and merged with the junctional bulge. To understand the physical forces behind the observations, we considered theoretically the mechanical energy of a membrane system consisting of a three-way tubular junction with emerging tubular arms connected to a membrane reservoir. The energy minimization predicted the membrane bulging, preferably, in the junction site as a result of both an increase in the luminal pressure and an abrupt rise of the membrane tension. We discuss the common physical background of the two phenomena.

## Introduction

Migrasomes are the recently discovered transient cell organelles whose biogenesis and evolution are tightly related to cell migration(1). Cells crawling on extracellular substrates leave behind retraction fibers (RFs), tens of microns long and about hundred-nanometer thick membrane nanotubules pulled out of the rear face of the cell. The retraction fibers are anchored to the substrate at their tips and in discrete locations along their lengths(2). Importantly, most of the retraction fibers are extensively branched with splitting sites represented by symmetric three-way tubular junctions(1). Migrasome formation and maturation are initiated by local swellings of retraction fibers into micron-scale spherical bulges. The retraction fibers degrade with time, which results in migrasome separation from the cell and transformation into extra-cellular vesicles (EVs).

Evidence is being accumulated of the multiple essential functions of migrasomes related to their ability to encapsulate and transport a large variety of cargoes. Migrasomes are involved in such processes as morphogenetic signaling(3), cell-cell communication(3, 4), inter-cellular transport of mRNA and proteins(4), removal from the cell of damaged mitochondria(5), spreading of diseases including cancer metastasis(6, 7) and the pathogenesis of brain injury(8).

Understanding the fundamentals of migrasome biogenesis and, importantly, the development of strategies for migrasome usage for therapeutic and diagnostic purposes require knowledge of the critical molecules and physical forces involved in the migrasome shaping(9). The established crucial molecules are cholesterol and the specific members of the tetraspanin (TSPAN) family of proteins(10) such as TSPAN4 and/or TSPAN7(9). A suggested mechanism of the migrasome shaping assumed formation within the RFs of macroscopic membrane domains enriched in TSPAN and cholesterol. A large mechanical stiffness of these domains was proposed to drive local swellings of the RF membrane(9).

A recent study provided further phenomenological insight into the mechanism of migrasome formation. It was shown in cell and biomimetic systems that migrasomes emerge in two steps(11). First, local vesicle-like swellings arise on the retraction fibers with a relatively weak change in the membrane composition of the swollen regions. While the swellings have been observed to form at different positions along the RFs, their preferable locations were the RF tips and junction sites. This is followed by the second step in which the swellings grow and mature into stable migrasomes through a strong enrichment in TSPAN(11). In case TSPAN is absent from the system, the swellings that emerged in the first step quickly disappear and the RF membrane recovers its initial configuration(11). This suggests that the major role of TSPAN is in the stabilization of the migrasome structure(9), while the initial local swellings of the RF membrane tubules can be driven by other and, possibly, unspecific factors such as mechanical forces.

The goal of the present study is to advance the understanding of the migrasome biogenesis by addressing the mechanisms of the first TSPAN-independent step of this process. Using the live cells and the biomimetic model system of membrane tethers pulled out of giant liposomes we demonstrate that the initial swellings can be produced by unspecific mechanical factors: an osmotically produced excess of intraluminal pressure or a rapidly generated excess of membrane tension. We demonstrate that the nascent migrasomes appear preferentially on tubular junctions while those formed on tubules tend to migrate into the junctions. In the absence of TSPAN, the emerged migrasomes disappear over time in both the cell RF and the biomimetic systems. Using the theory of membrane elasticity, we elaborate on the criteria of the swelling generation on the tubules and tubular junctions upon sudden increases of the pressure or the membrane tension. We demonstrate that the junctions are mechanically more susceptible to swelling than the tubules so that the nascent migrasome formation at the tubular branching sites is kinetically preferable and energetically favorable.

## Experimental Results

### Increase of luminal pressure in retraction fibers generates swellings preferentially in junctions

First, we aimed to test whether luminal pressure can trigger swelling in retraction fibers of live migrating cells. For this purpose, we exposed RF-forming culture cells to a hypotonic medium. The idea behind this experiment was that the excess of osmotic concentration in the RF lumen generates a water influx through the RF membrane into the RF lumen, which in turn drives a water flow along the RF into the cell body, the latter serving as a reservoir with slowly changing pressure. This water flow generates within the RF a hydrodynamic pressure, which pushes on the RF membrane and represents the RF luminal pressure. The flow ceases after a relatively long period of time needed for the osmotic equilibration between the system interior and the outside medium.

The cell exposure to a hypotonic solution stimulated the formation of swellings on the initially smooth RFs (Fig.1). The swellings gradually developed within a time scale of a few tens of seconds (Fig.1A). After longer time periods of about 3 – 5 minutes the swellings disappeared. The typical swelling size was on a micron scale and slightly depended on the external osmolarity (Fig. 1 B,C). Importantly, more than 80% of the swellings formed on the three-way junctions (Fig. 1 D,E). Note that the junctions appeared either as regular symmetric three-way vertices with 120-degree angles between the rays or as irregular vertices in which one ray was hardly recognizable, apparently, because of its shortness.

**Figure 1.**
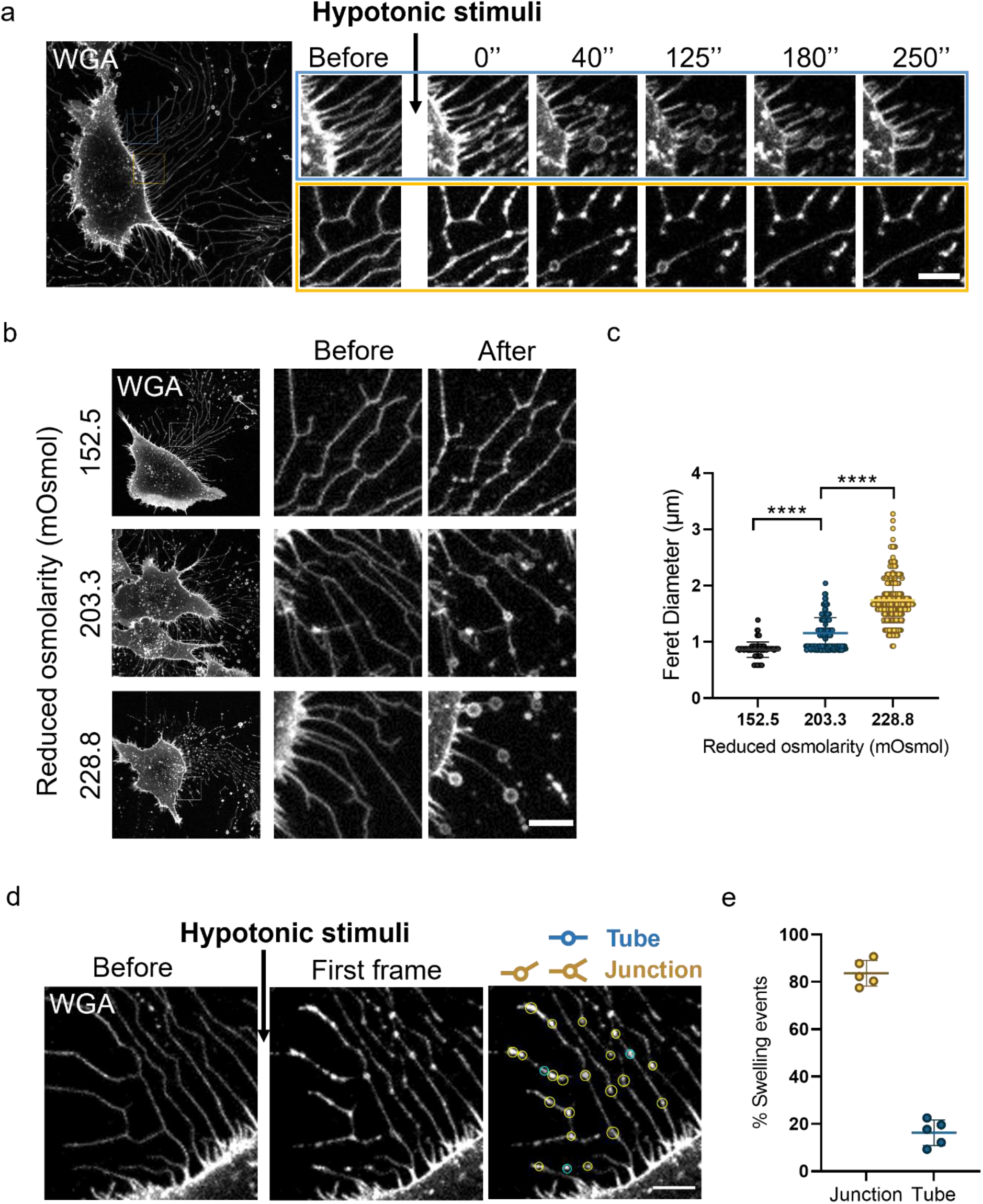
Swellings formation in retraction fibers by cell exposure to hypotonic medium.L929 cells pre-stained with WGA488 in DPBS were used for all experiments in this figure. (A) Image series showing the progress of hypotonic induced swellings on retraction fibers. Scale bar, 5 μm. (B) Representative confocal images of cells before and after treatment with hypotonic buffers. The reduced extracellular osmolality was labeled at the right side of each panel. Scale bar, 5 μm. (C) Statistical analysis of the diameter of swellings in (B). Data acquired from 3 independent experiments were pooled for analysis. A total of 116, 252 or 219 swelling events were analyzed respectively. P values were calculated using a two-tailed unpaired nonparametric test (MannWhitney test). P value<0.05 was considered statistically significant. **** P<0.0001. (D) Representative confocal images of a cell before and after hypotonic stimulation. Cells were treated with 25% DPBS (228.8 mOsmol osmolality was reduced). Swelling events in tubes or junctions were highlighted by blue or yellow circles respectively. (E) Statistical analysis of the site of swellings in (D). Data acquired from 3 independent experiments were pooled for analysis. A total of 273 swelling events in 5 cells were analyzed. Scale bar, 5 μm.

### Abrupt increase in membrane tension generates swellings preferentially in tubular junctions

To test the effects of fast increments of membrane tension on the morphology of branched retraction fibers (RFs) we used a biomimetic system of three tubules connected by a three-way junction (Fig.2A). To create this system, Giant plasma membrane vesicles (GPMVs) were generated from HEK293T cells (see Methods). Next, the vesicles were added into a custom-made chamber mounted on the stage of a correlated optical tweezers confocal fluorescence microscope. A micropipette aspiration setup was incorporated into the tweezers-confocal system. The GPMV membrane was subjected to tension by micropipette aspiration such that the tension values were determined by the controlled suction pressure (see Methods). Two optically trapped polystyrene beads were pushed towards an aspirated GPMV and two membrane nanotubes were pulled out of the vesicle (Figs. 2A). The distance between the tubules was reduced by moving one of them towards the other until the tubules coalesced and formed a 3-way junction (Fig. 2A)(12).

**Figure 2.**
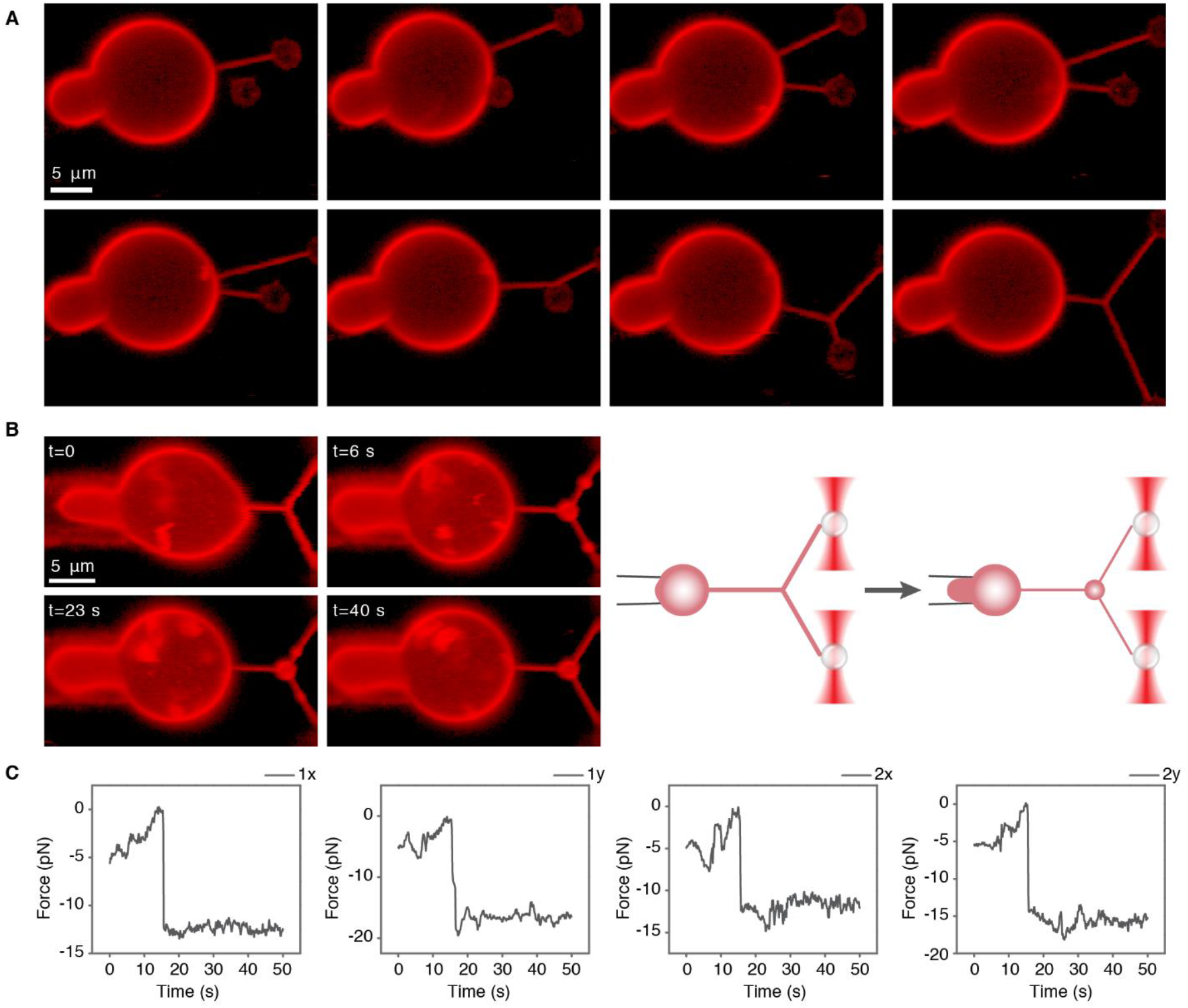
Swellings formation by abrupt tension increase in the biomimetic system. (A) 3-tube junction formation. Confocal microscopy images of 3-tube junction formation from aspirated GPMVs dyed with DiI-C12. Two membrane tubes were pulled from the vesicle. Next, the upper tube was elongated and brought towards the other tube until the tubes coalesced. (B) Swelling formation by tension jump. Time-lapse confocal microscopy images of the tension jump assay. First, the suction pressure was reduced to zero (t=0), then the suction pressure was increased instantaneously to 0.39 mbar (t=6 s), corresponding to 0.074 mN/m membrane tension, leading to swelling formation. Small swellings upstream of the junction merged with the junction (t=40s). A video of this process can be seen in supplementary figure 1. This experiment was performed 17 times on 11 different GPMVs in 5 independent experiments. A schematic illustration is shown on the right. (C) Force plots of the tension jump presented in B. The left panels correspond to the x and y force components measured from the bead1 displacement from optical trap1 and the right panels correspond to the forces measured from the bead2 displacement from optical trap2. The fluorescence intensity of all the images is presented in log scale.

First, the tubular system was generated with a relatively low membrane tension upon a minimum suction pressure (Fig. 2B, t=0). Then, the suction pressure was instantaneously increased to values in the range of 0.2 - 0.44 mbar corresponding to the GPMV membrane tension in the range of 0.04 – 0.1 mN/m. This resulted in the formation within the tubular system of membrane swellings reminiscent of migrasomes ((Fig.2B) and supplementary video S1). Immediately upon an abrupt increase in the membrane tension, swellings appeared in the junction and in the tubules upstream of the junction (Fig. 2B at *t*= 6*s*). The tension increase applied by changing the aspiration pressure could also be observed as a step change in the force needed to hold the tubes (Fig. 2C). At later stages, the tubule’s swellings moved towards the junction (Fig. 2B at *t*=23*s*) and, finally, merged with the junctional swelling (Fig. 2B at *t*=40*s*).

### Modeling the mechanics of swelling generation

To understand the physical basis for the above phenomena of the local membrane swellings we developed a theoretical model. The common feature of the two observed occurrences of the swelling generation is the mechanical origin of the driving forces, the luminal pressure in the branched retraction fibers and the membrane tension in the biomimetic system. Therefore, the two systems must share the physical principle underlying the local swelling - the tendency to minimize mechanical energy. In the following, we formulate a joint thermodynamic model describing both an element of a branched retraction fiber consisting of one 3-way junction with three tubular arms and the biomimetic system of three membrane tubules connected by a junction. We analyze the model predictions separately for each system.

### Model

We consider a membrane system consisting of three membrane tubular arms emerging out of a 3- way junction and symmetrically oriented with 120° angles between the tubular axes (Fig. 3A). The distance from the junction center to the arm end is denoted by *L*. The ends of the tubular arms are merged with a reservoir of the lipid material building the system’s membrane and of the aqueous medium filling the system’s luminal volume. The cross-section of the boundary line along which a tubular arm is connected to the reservoir is assumed to be a circle of radius *R*.

**Figure 3.**
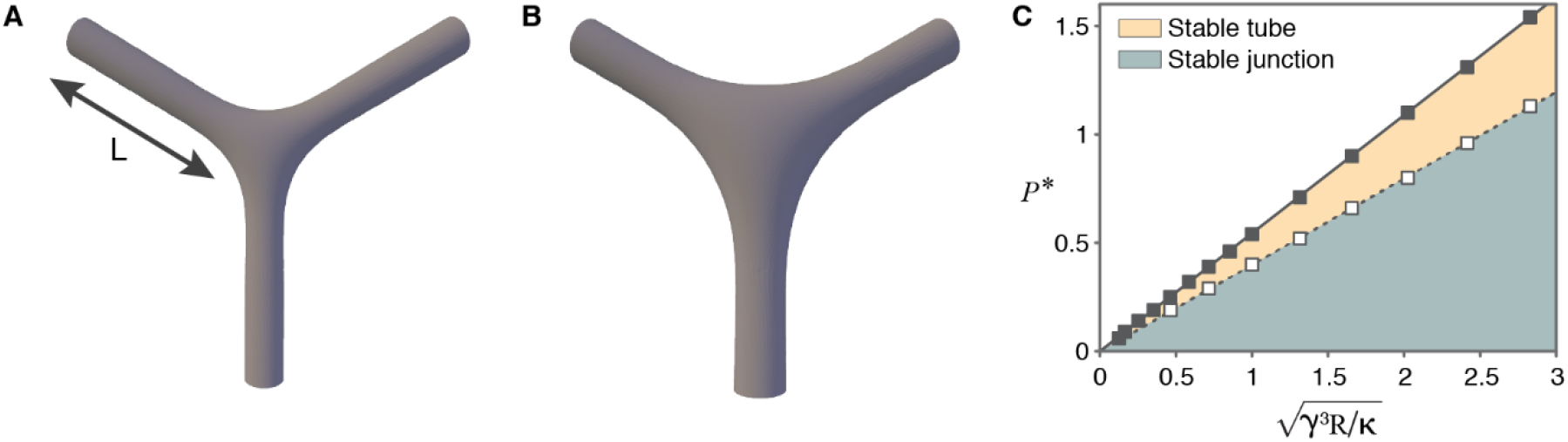
Modeling junction swelling and instability by increasing pressure. The tubules and junctions were computed by Brakke’s Surface Evolver (based on the method of finite elements) for a membrane system characterized by a bending rigidity, *κ*, a tension *γ*_*R*_ and apositive luminal pressure, *P*. (A) A three-way junction connecting three tubules with vanishing intraluminal pressure, *P*=0. (B) The same as (A) but with positive pressure *P* > 0. The elastic parameter is 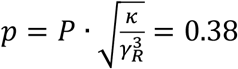, the length of the tubules is 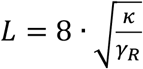. (C) Phase diagram for the stability of the junctions and tubules. The straight lines represent the phase boundaries and correspond to the critical pressure for junctions and tubules.

The reservoir imitates the GPMV for the biomimetic system and the cell body for the RF element. In reality, in the biomimetic system, the end of only one tubular arm is in contact with the reservoir, while the two others are held by the pulling forces produced by the laser traps whereas for the RF element, the ends of all three tubular arms are subjected to pulling forces. Yet, for both the biomimetic and RF systems due to the self-connectivity and fluidity of the membrane and of the luminal liquid, all tubular arms eventually exchange the membrane material and the aqueous content with the reservoir and, therefore, can be modelled as being effectively directly connected to the reservoir.

We consider the reservoir membrane to be flat and subjected to lateral tension, *γ*_*R*_, and the reservoir liquid content to be under a hydrostatic pressure exceeding the pressure in the outside medium by *P*. In the following, we refer to *P* as the pressure and to *γ*_*R*_ as the tension, for brevity.

We define the system energy, *F*, as a thermodynamic work of the system generation out of the reservoir, which can be seen as consisting of three sub-processes each associated with an energy cost: pulling out of the reservoir the membrane area, *A*, and the volume of the aqueous medium, *V*, characterizing the system, and bending the initially flat pulled membrane into the shape of the system. The related thermodynamic work is

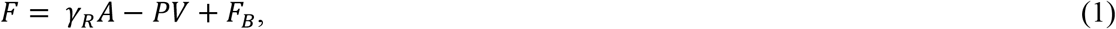

where the first and second contributions describe the work of pulling, respectively, the membrane area and aqueous volume out of the reservoir, while the third contribution represents the energy of the membrane bending. We treat the bending energy, *F*_*B*_, according to Helfrich model(13). We quantify the local membrane shape at each point of the membrane surface by the total curvature, *J*, which is the sum of the two principal curvatures, and consider the area density of the local bending energy, *f*_*B*_, to be given by

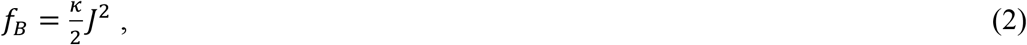

where *κ* is the membrane bending modulus(13). The total membrane bending energy is given by

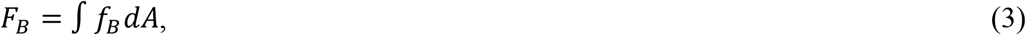

where the integration is performed over the entire area of the system’s membrane.

By using (Eq.2) we assume the membrane to have a vanishing spontaneous curvature(13), which implies that the two membrane monolayers have identical elastic properties providing the membrane with up-down symmetry. In addition, we do not consider the energy contributions related to the membrane Gaussian curvature(13) since the phenomenon of local membrane swelling, which we are interested in, does not affect the membrane topological genus so that the Gaussian curvature energy remains constant according to the Gauss-Bonnet theorem(14).

Analytical minimization of the energy (Eq.1) results in a non-linear differential equation for the membrane curvature referred to as the shape equation(15, 16). In the particular case of a cylindrical membrane, the shape equation acquires a simple form,

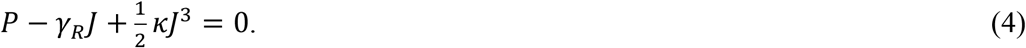

According to (Eq.4), the membrane configurations have an intrinsic length scale 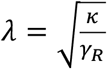, which represents the typical relaxation length of the membrane curvature. In the following we assume the length of the tubular arms to be sufficiently large, *L* ≫ λ, so that the ends of the tubules are, practically, not influenced by the junction.

#### Way of calculations

Generally, the configuration of the system could be obtained by solving the shape equation. Yet, because of the threefold symmetry of the tubular junction and the possible deviations of the tubular arms’ shapes from those of smooth cylinders, we cannot use the simple form of the shape equation (Eq.4) while solving the shape equation in its general form (15, 16) appears too complicated. Therefore, we numerically minimized the total energy of the system (Eq.1) by using Brakke’s “Surface Evolver” (17).

In the following, we use the model to predict the energetically favorable configurations of the system for two experimental conditions described in the previous sections: the system inflation by increasing luminal pressure and exposure of the system’s membrane to an abrupt increase of the membrane tension.

We start with computing the system’s initial configuration corresponding to a tension, *γ*_*RO*_, and a vanishing pressure, *P*=0. The energy minimization results in the configuration presented in (Fig. 3A). The tubular arms far enough from the junction asymptotically acquire the shape of a smooth circular cylinder with a cross-section radius proportional to the relaxation length, 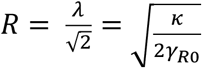, which automatically matches the boundary condition at the entry to the reservoir. The energy of this configuration will be denoted by *F*_0_.

### Junction swelling by increasing pressure

Here we consider the transformation of the system configuration resulting from an increase of the pressure to a positive value, *P* > 0, upon keeping constant the reservoir membrane tension, *γ*_*R*_, equal to its initial value, *γ*_*RO*_. Based on the shape equation(15, 16) for the general case and (Eq.4) for a cylindrical shape, the membrane shapes depend on a dimensionless parameter

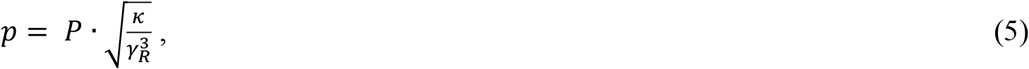

referred below to as the elastic parameter.

Our computation demonstrates the existence of a critical elastic pressure, *P*^∗^, corresponding to the elastic parameter, *p*^∗^ ≈ 0.4, which separates two distinct regimes of the evolution of the system configuration.

As long as *p* remains smaller than the critical values, *p* < *p*^∗^, the pressure growth results in a monotonic expansion of the junction and inflation of the tubular arms (Fig. 3B). At distances from the junction’s center larger than the relaxation length λ, the tubules asymptotically acquire a shape of a circular cylinder (Fig.3B) whose cross-sectional radius is related to the tension and the pressure through the solution of the equilibrium equation for a cylinder (Eq.4).

The critical pressure, *P*^∗^, corresponding to *p*^∗^, is presented as a function of tension in (Fig.3C).

Once the pressure exceeds the critical value so that, *p* ≥ *p*^∗^, the junction loses a stable shape and tends to unlimitedly inflate. This instability corresponds to a swelling formation in the junction. The swelling stops growing if the pressure drops during the inflation.

It is instructive to compare the condition of the junction instability with that of a homogeneous cylindrical tubule. A membrane cylinder becomes unstable and tends to unlimitedly inflate if the equation of equilibrium (Eq.4) does not have a positive solution for the tubule’s radius, *R*. It can be shown analytically that this happens if the elastic parameter of the cylinder exceeds the critical value 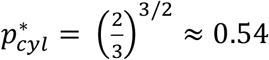, which is larger than the critical pressure of the junction *p*^∗^ ≈ 0.4. The critical pressure of a cylinder is presented in (Fig.3C) aside of those of a junction. Hence, junctions are more susceptible than tubules to the loss of stability and swelling by the luminal pressure. This explains our numerical result according to which in the considered system of a junction with emerging tubular arms, the junction is the first to lose the equilibrium configuration and start swelling upon an increasing pressure.

This conclusion of the model explains the above experimental observation (Fig.1) of a preferential junction swelling in the system of retraction fibers exposed to increasing luminal pressure.

### Junction swelling by abrupt increase of the membrane tension

We consider the effect on the system configuration of a change of the tension from *γ*_*RO*_ to *γ*_*R*_.

The character of the system shape transformation depends on the relationship between the pace of the tension variation and that of the volume exchange between the system lumen and the reservoir, which is mediated by the viscous flow of the luminal liquid. If the tension alteration is sufficiently slow so that the volume is concomitantly redistributed, the pressure within the system remains constant and equal to that of the reservoir, *P*=0. In this case, the system configuration remains geometrically similar to that of the initial state (Fig. 3A), while all the system dimensions rescale with the ratio of the relaxation lengths before and after the tension variation,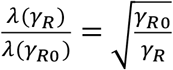

Hence, in this regime no swelling is predicted to develop in the system.

Here we analyze a case of an abrupt change of the tension during which the volume exchange between the system and the reservoir does not occur because of kinetic limitations of the viscous flow. Hence, we assume that within the short time of the tension variation the total volume of the system remains constant, *V*=const, but subject to local short-range redistributions within the system itself. Consequently, within the same time span, the luminal pressure of the system can deviate from that of the reservoir.

#### Pearling of junction

Our numerical computations show an abrupt application of an elevated tension to result in a narrowing of the tubular arms and inflation of the junction, which is accompanied by a redistribution of the luminal volume from the tubules to the junction. Another consequence of the tension application is the generation within the system of a luminal pressure, *P*, which overcomes the vanishing pressure in the reservoir because of the restriction of the volume exchange.

According to our results, there are two regimes of the change of the system conformation. If the applied tension value, *γ*_*R*_, is relatively close to *γ*_*RO*_, the system configuration (Fig. 4A) remains qualitatively similar to the initial one (Fig. 3A). The tubular arms become narrower while the junction inflates but its projection on the system plane retains the shape of a triangle with concave edges (Fig. 4A). The amount of the redistributed volume and, hence the extents of the tubule narrowing and of the junction inflation are set by the values of the initial, *γ*_*RO*_, and the abruptly applied, *γ*_*R*_, tensions, and of the total length of the tubular arms.

**Figure 4.**
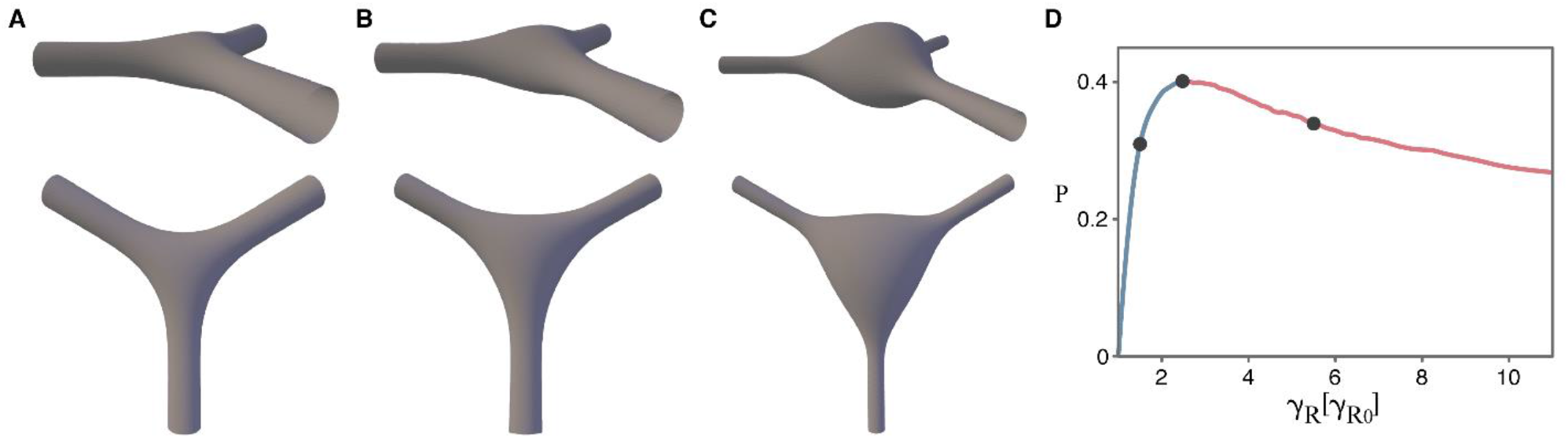
Modeling pearling of junctions by an abrupt increase of tension. (A-C) A junction connecting three tubules with fixed volume and increasing tensions. The junction is initially in mechanical equilibrium under an initial tension *γ*_*RO*_. (A) A small increase in tension to *γ*_*R*=_1.5*γ*_*RO*_ results in a modest expansion of the junction, as seen from the top view (bottom panel), while the contours of the junction sides remain concave. (B) The generated tension has a critical value *γ*_*R*=_2.5*γ*_*RO*_ corresponding to the critical values of the elastic parameter, *p*^∗^ ≅ 0.4. (C) The generated tension, *γ*_*R*=_7*γ*_*RO*_, exceeds the critical value resulting in pearling of the junction. The contours of the junction sides are convex while overall shape of is more spherical. The length of tubules is 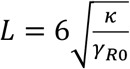 and the volume of the system is 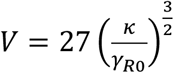. (D) The elastic parameter as function of the tension. The blue and red line colors indicate, respectively the pre- and post-critical regimes of the junction swelling.

If the applied tension exceeds a critical value, 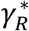, the character of the system’s evolution changes. The shape of the junction projection turns to a triangle with convex edges (Fig. 4C). In this regime, which is referred below to as the junction pearling, the junction shapes become progressively similar to a sphere.

To understand the origin of the critical tension, 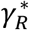, triggering the onset of the junction pearling, we considered the elastic parameter, *p*, introduced in the previous section for the pressure-induced swelling (Eq.5). In the present case, the pressure in (Eq.5) is not set independently but rather depends on the tension, *P*(*γ*_*R*_), according to the results of our numerical computations. The dependence of *p* on the ratio between the applied and the initial tensions, *γ*_*R*_/*γ*_*RO*_, has a non-monotonic character with a maximal value, *p*^∗^ ≈ 0.4 (Fig.4D). This maximal *p*^∗^ is the critical value of the elastic parameter corresponding to the junction pearling transition, which is represented in (Fig. 4B). The obtained critical *p*^∗^is the same as that found above for onset of the junction instability with respect to swelling upon increasing pressure and constant tension. The critical tension, 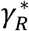, can be found by solving the equation

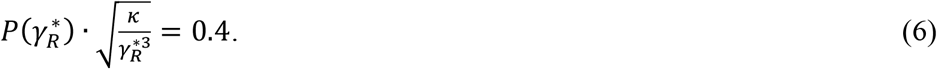

Note that the numerically computed function, *P*(*γ*_*R*_) depends on the initial tension, *γ*_*RO*_, and the lengths of the tubular arms, *L*.

The obtained criterion of the pearling transition in the junction can be compared with that of pearling of cylindrical membrane tubules of constant volume subject to membrane tension(18). Our numerical computations showed that the onset of the tension-driven formation of a sphere-like bulge on a tubule with constant volume is described by the critical value of the elastic parameter very close to, *p*^∗^ ≈ 0.54. The same value of the elastic parameter was found above for the onset of instability of tubules inflated by a luminal pressure. Hence, the three-way tubular junctions are more susceptible than tubules to the pearling transition driven by an abrupt tension application.

#### Comparison of different regimes of pearling

Since our experiments with abrupt application of tension resulted in a transient coexistence of swellings in the junction with those in the tubular arms (Fig .2B), we assumed that if the applied tension exceeds the critical pearling tensions of both the junction and the cylindrical tubes, such coexistence may represent a state of local equilibrium, while the sole pearling of the junction would be the state of the global equilibrium.

To verify this hypothesis, we considered three types of scenarios: (i) pearling of one tubular arm only, (ii) pearling of the junction only, and (iii) a combination of the junction and the tubular arm pearling. We numerically found that all these configurations correspond to minima of the system energy, and therefore indeed represent the local equilibrium states. The three equilibrium configurations differ in their energies related to the energy of the initial state, which are presented in (Fig. 5A). According to these results (Fig.5A), the pearling of junction only is the most energetically favorable configuration of the system, which, hence, represents the state of the global energy minimum.

**Figure 5.**
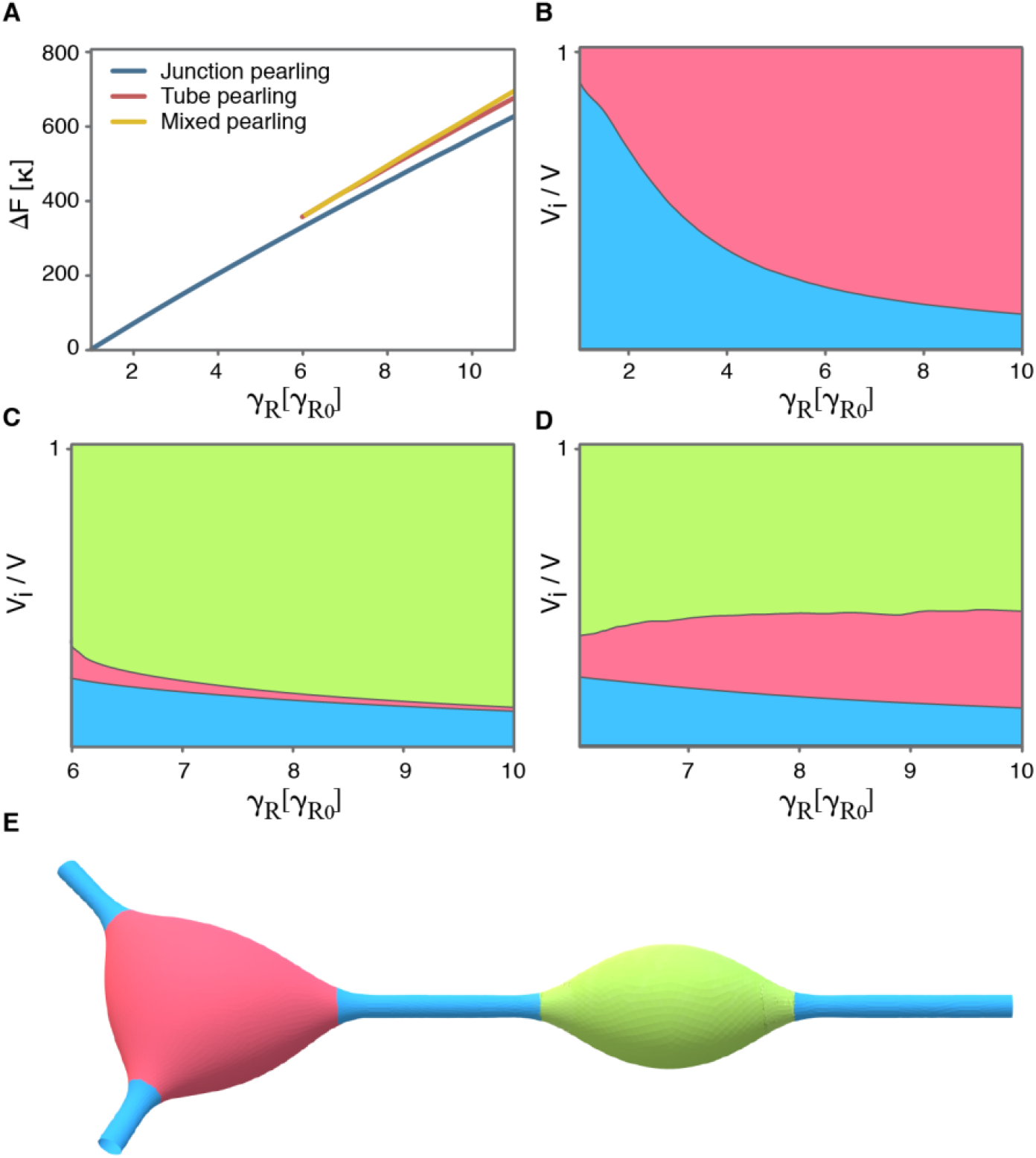
Different regimes of pearling in the system. Energy minimization reveals three regimes of pearling each resulting in a state of local minimum of the system energy: (i) pearling of the junction (blue curve), (ii) pearling of the tubular arm (orange curve), and (iii) combination of pearling in the junction and the tubular arm(yellow curve). (A) The energies, of the three states as functions of the tension, *γ*_*R*_. (B-D) The volume distribution between different compartments in each regime. The fractional volume of each compartment is described by a color, as illustrated in (E): *V*_*junction*_ in red, *V*_*tubules*_ in blue, and *V*_*tubule*−*pearling*_ in green. (E) Example of a type (iii) state. The length of a tubular arm is 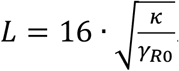.

The three equilibrium configurations differ also in their volume distribution between the different sub-systems, namely, the junction, the tubular regions of the arms, and arm’s bulge. The volume fractions of the sub-systems for the three scenarios are shown in diagrams (Fig.5 B,C,D) as functions of the applied tension. A particular case of a system configuration in the coexistence state is presented in (Fig. 5E).

The obtained relationships between the energies of the three scenarios (Fig. 5A) explain the experimentally observed time evolution of the system in which the swellings initially formed on the tubular arms ultimately fuse with the junctional swelling.

On sufficiently long time scales enabling full exchange on the luminal fluid between the system and the reservoir, the constant volume condition resulting in the pearling transition is relaxed so that the system swellings are predicted to vanish, which agrees with our observations.

## Discussion

Migrasomes are micron-scale vesicle-like organelles emerging on cell retraction fibers (RFs) during cell crawling on external substrates(1). The preferred location of migrasomes are the sites of 3-way junctions between the tubular regions of branched RFs(1). The migrasome formation was shown to occur in two steps: the first, apparently, protein-independent step of the initial local RF swelling into micron-large spherical bulges, and the second step of the bulge maturation and stabilization by tetraspanin (TSPAN) proteins. While the physical factors underlying the second step were studied(9, 11), the forces responsible for the first step have not been addressed.

Here we proposed that the first step of migrasome formation can be mediated by the generic mechanical factors, an excess of pressure in the RF lumen with respect to that of the outside medium or an abrupt jump of the RF membrane tension. We substantiated this proposal experimentally and theoretically. We exposed live cells lacking TSPANs and having well-developed systems of retraction fibers to a hypoosmotic medium and observed the transient formation of membrane bulges preferentially in the junction sites of RF tubular branches. In a biomimetic system of three membrane tubules connected by a junction, we demonstrated that an abrupt tension jump produces a migrasome-like swelling with a preferred location in the junction site. Finally, we theoretically analyzed the energetically favorable configurations adopted by a system of three membrane tubules connected by a three-way junction upon increase of the intra-tubular pressure or an abrupt increase of the membrane tension. In both cases the computations predicted formation of localized membrane swellings, whose location in the junction sites is energetically more favorable than in the tubular regions. Altogether, we presented compelling evidence for a mechanical origin of the forces initiating the migrasome formation in the three-way junction sites of branched retraction fibers.

In a general perspective, the formation of membrane swellings referred to as the pearling transition or peristaltic instability has been extensively studied for cylindrical membrane tubules and appears to be a rather ubiquitous phenomenon typical for both lipid vesicles and cell membranes(19, 20). Phenomenologically, the transition consisting in the stability loss of the initially smooth tubular shapes and formation of chains of bead-like membrane bulges joined by cylindrical connections looked qualitatively very similar for membrane tubules of different and completely unrelated origins. Yet despite the resemblance of the membrane configurations, the physical factors driving the transitions varied for different systems. The pearling transition was observed to result from a longitudinal stretching of tubule-like vesicles upon conservation of the tubule’s volume(18), an osmotic swelling of axons by dilution of the external medium(21), generation of a large intrinsic curvature of membranes by membrane-bound or inserted molecules(20, 22), depolymerization of intracellular actin(19, 23). The pearling transition resulting from the vesicle stretching was proposed to be analogous to tension-driven Rayleigh instability opposed by the membrane resistance to bending rigidity(18), hydrodynamic drug(24), rigidity of the submembrane actin shell (19), gel-like dynamics of the cytoskeleton and the membrane-cytoskeleton coupling(21).

The phenomena studied here both experimentally and theoretically can be considered as a generalization of the pearling transition to systems of branched membrane tubules containing junctions. The essence of our finding is that tubular junctions are more susceptible to the pearling transition than tubules. This is true for the two kinds of the pearling-driving conditions we explored, an increase of the luminal pressure and an abrupt increase of the membrane tension. Our theory predicted the transition in the junction to occur at lower pressure and lower tension. This conclusion is supported by our observations of generation of the migrasome-like swellings, preferably, in the junctions of the retraction fibers, and of the tendency of the swellings formed on the tubular arms of the biomimetic system to merge with the swelling in the junction.

It is important to emphasize that in cell membranes the purely physical picture described and analyzed here must be modulated by specific biological factors. A recent work demonstrated an essential role of PIP5K1A, a PI4P kinase which converts PI4P into PI(4,5)P_2_ (25), and of sphingomyelin synthase 2 (SMS2) in migrasome biogenesis (26). Foci of SMS2 protein initially formed at the cell leading edge move to the retraction fibers during the cell migration and serve as the sites of migrasome nucleation. According to the proposed model, in case a SMS2 foci is formed within a tubular region of a retraction fiber, the resulting nascent migrasome must be moved into a junction by mechanical forces. Another possibility is that such nucleation site has a lower chance to develop into a migrasome compared to a SMS2 foci located at three-way junctions.

A possible biological rationale for migrasome formation in the junctions rather than the linear tubular regions of branched retraction fibers is a kinetic facilitation of the migrasome loading with luminal and membrane cargos.

## Materials and Methods

### Hypotonic stimuli applied to retraction fibers

#### Cell culture

L929 cells (for cell imaging) or HEK293T cells (for biomimetic system) were cultured at 37 °C, 5% CO_2_ in DMEM (Gibco) high-glucose medium supplemented with 10% fetal bovine serum (Biological Industries), 1% glutamax (Gibco) and 1% penicillin–streptomycin.

### Live cell imaging

Cells were seeded in a glass-bottom dish (Cellvis, D35C4-20-1.5-N) and allowed to grow for 14 -18 hours. The dish was pre-coated with 10 μg /ml fibronectin (Sigma, F0895) at 37 °C to enhance cell attachment. Before imaging, culture medium was gently replaced with a staining solution (2 μg /ml WGA Alexa Fluor™ 488 conjugate (Invitrogen) in DPBS) and cells we incubated for 10 min to achieve bright labeling of plasma membrane. Cells were then transferred to a Nikon A1 laser scanning confocal microscope equipped with a live cell system. For all experiment, a 5 min 3D time-lapse series was collected to fully capture the progress of swelling. Images were captured with 8s interval, at each time point, a Z stack containing 3 slices with 500 nm step size was recorded.

Real-time hypotonic stimulation during time-lapse imaging was achieved by using a home-made buffer-exchange equipment (refer to the eMigrasome paper under preparation).

### Giant plasma membrane vesicles (GPMVs) formation

GPMVs were produced according to a published protocol(27). Briefly, HEK293T cells were stained with DiI-C12 (Invitrogen) membrane dye, washed with GPMV buffer (10 mM HEPES, 150 mM NaCl, 2 mM CaCl2, pH 7.4) twice, and incubated with 1 mL of GPMV buffer containing 1.9 mM DTT (Sigma) and 27.6 mM formaldehyde (Sigma). Secreted GPMVs were then collected and isolated from the cells and immediately used for optical trapping experiments.

### 3-tube junction formation form aspirated GPMVs

The experiments were performed using a C-trap ® confocal fluorescence optical tweezers setup (LUMICKS) made of an inverted microscope based on a water-immersion objective (NA 1.2) together with a condenser top lens placed above the flow cell. The optical traps are generated by splitting a 10W 1064-nm laser into two orthogonally polarized, independently steerable optical traps. To steer the two traps, one coarse-positioning piezo stepper mirror and one accurate piezo mirror were used. Optical traps were used to capture polystyrene microbeads. The displacement of the trapped beads from the center of the trap was measured and converted into a force signal by back-focal plane interferometry of the condenser lens using two position-sensitive detectors. The samples were illuminated by a bright field 850-nm LED and imaged in transmission onto a metal-oxide semiconductor (CMOS) camera. **Confocal fluorescence microscopy:** The C-Trap uses a 3 color, fiber-coupled laser with wavelengths 488, 561 and 638 nm for fluorescence excitation. Scanning was done using a fast tip/tilt piezo mirror. For confocal detection, the emitted fluorescence was descanned, separated from the excitation by a dichroic mirror, and filtered using an emission filter (Blue: 500-550 nm, Green: 575-625 nm and Red: 650-750 nm). Photons were counted using fiber-coupled single-photon counting modules. The multimode fibers serve as pinholes providing background rejection. Experimental chamber: PDMS walls were placed on the bottom cover slip (Bar Naor) and mounted onto an automated XY-stage. The GPMVs sample was added to the chamber and a few drops of oil were put on the sample surface to prevent evaporation. **Micropipette aspiration:** A micropipette aspiration setup including micromanipulator (Sensapex) holding a micropipette with diameter of 5 μm (Biological industries) connected to a Fluigent EZ-25 pump was integrated to our optical tweezer instrument. Before and after each experiment, the zero-suction pressure was found by aspirating a polystyrene bead (3.43 μm, Spherotech) into the pipette and reducing the suction pressure until the bead stopped moving. **Tubular 3-way junction formation**: In order to pull a membrane tube, an optically trapped bead was brought in contact with the GPMV for about a minute, and then moved away from the vesicle(28).

First, two membrane tubes were pulled from aspirated GPMVs using two optically trapped beads (12). The distance between the tubes was reduced by moving one of the tubes towards the other, until they coalesced and formed 3-membrane tube junction. To form swellings, the suction pressure was reduced to minimum. Then, we instantaneously increased the suction pressure to values in the range of 0.2-0.44 mbar (correspond to 4-10×10^−5^ N/m membrane tension). For the confocal imaging 532 nm laser was used for DiI-C12 excitation with emission detected in the green channel.

## Acknowledgements

MMK was supported by Israel Science Foundation (Grant No. 1994/22) and holds Joseph Klafter Chair in Biophysics. Raya Sorkin acknowledges support by the Israel Science Foundation (Grant No. 1289/20), and the NSF-BSF (Grant No. 2021793) and holds the Raymond and Beverly Sackler Career Development Chair for Young Faculty. Co-Funded by the European Union (ERC *ReMembrane 101077502*). Li Yu by the National Natural was supported by Science Foundation of China to Li Yu (grant no. 32030023 and 92054301). Views and opinions expressed are however those of the authors only and do not necessarily reflect those of the European Union or the European Research Council Executive Agency. Neither the European Union nor the granting authority can be held responsible for them.

## Supplementary Video legend

3-tube junction formation followed by swelling formation. A video of formation of 3-tube junction pulled from aspirated GPMV dyed with DiI-C12. Next the aspiration pressure was reduced to zero and then it was increased instantaneously to 0.39 mbar (corresponding to 0.074 mN/m). The video was composed of confocal fluorescence microscopy images of the experiment shown in Figure 1b. The fluorescence intensity is presented in logarithmic scale.

