## Supplementary figures and images for "Migrasome formation is initiated preferentially in tubular junctions by alternations of membrane tension or intracellular pressure"

### supplementary movie

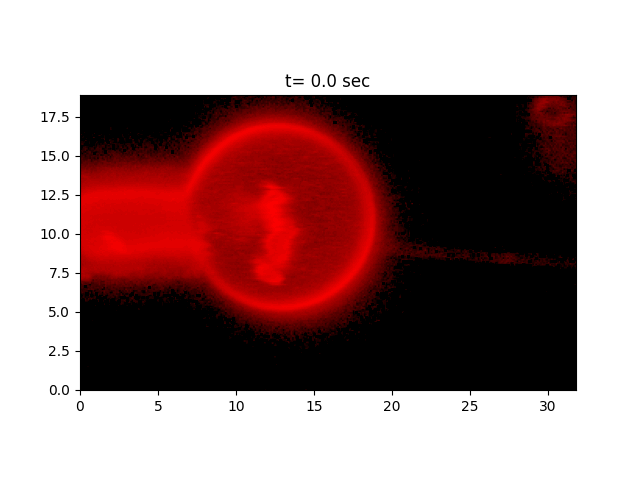
